# New cinnamid and rosmarinic acid derived compounds biosynthesized in *Escherichia coli* as *Leishmania amazonensis* arginase inhibitors

**DOI:** 10.1101/2020.10.30.362111

**Authors:** Julio Abel Alfredo dos Santos Simone Come, Yibin Zhuang, TianZhen Li, Simone Brogi, Sandra Gemma, Tao Liu, Edson Roberto da Silva

## Abstract

Arginase is a metalloenzyme that plays a central role in *Leishmania* infection. Previously, rosmarinic and caffeic acids were described as antileishmanial and as a *Leishmania amazonensis* arginase inhibitor and now, we describe the inhibition of arginase in *L. amazonensis* by rosmarinic acid analogs (**1-7**) and new caffeic acids derived amides (**8**-**10**). Caffeic acid esters and amides were produced by means of the engineered synthesis in *E. coli* and tested against *L. amazonensis* arginase. New amides (**8**-**10**) were biosynthesized in *E. coli* cultured with 2 mM of different combinations of feeding substrates. The most potent arginase inhibitors showed Ki(s) between 2 - 5.7 μM. Compounds **2-4** and **7** inhibited L-ARG through a noncompetitive mechanism, and **9** showed a competitive inhibition. By applying an *in silico* protocol we determined the binding mode of compound **9**. The competitive inhibitor of L-ARG targets key residues within the binding site of the enzyme establishing a metal coordination bond with the metal ions as well as a series of hydrophobic and polar contacts supporting its micromolar inhibition of L-ARG. These results highlight that the dihydroxycinnamic-derived compounds can be used as the basis for the development of new drugs using a powerful tool based on the biosynthesis of arginase inhibitors.

## 1. Introduction

Leishmaniasis is a neglected tropical disease with a public health importance. While cutaneous leishmaniasis has affected 1 million people in the last 5 years, if untreated, visceral leishmaniasis results in over 20,000 deaths each year.^[1]^ There is a lack of effective drugs for leishmaniasis treatment, and the few available therapeutic options present high toxicity associated with treatment resistance.

Evolution of the leishmaniasis depends on the balance among Th1 cytokines (which triggers the activation of L-arginine metabolic pathways for nitric oxide (NO) production, responsible for killing the parasite) and Th2 cytokines, which determine the induction of host arginase and enhance the *Leishmania* infection.^[2]^ Arginase is an enzyme that catalyzes the hydrolysis of L-arginine to L-ornithine and urea. *Leishmania* is auxotrophic for polyamines (putrescine, spermidine, and spermine) that is important for the infection.^[3,4]^ Polyamines are substrates for the production of trypanothione, an antioxidant that neutralizes reactive oxygen species (ROS) and NO.^[5,6]^ Arginase activity deprives nitric oxide synthase 2 (NOS2) of the substrate, which decreases NO biosynthesis in the host defense cells, and provides polyamines, which can increase parasite proliferation.^[2]^

Due to the relevant action in the L-arginine metabolism pathway, arginase could be an important target for leishmaniasis treatment, and arginase inhibitors can reduce the parasite burden in *Leishmania*-infected BALB/c mice.^[7]^ We previously identified compounds with catechol groups^[8][9][10]^ and thiosemicarbazide^[11]^ as crucial *L. amazonensis* arginase (L-ARG) inhibitors. The compound verbascoside, a major constituent of the medicinal plant *Stachytarpheta cayennensis* that is used to treat leishmaniasis in Brazil^[12]^ and malaria in Peru^[13]^, inhibits L-ARG and it is active against both promastigote and intracellular amastigotes of the parasite.^[14, 15]^ Others natural dihydroxycinnamic compounds (caffeic acid, chlorogenic acid, and rosmarinic acid) exhibited *in vitro* and *in vivo* activity against *L. amazonensis*,^[16, 17]^ also showing inhibitory profile against L-ARG.^[18]^ Based on the antileishmanial activity of dihydroxycinnamic compounds,^[17]^ and L-ARG inhibition, we tested eight compounds previously produced via engineered *E. coli*.^[19, 20]^ In addition, we synthesized six new amides containing the 3,4-dihydroxycinnamic moiety and we tested them against L-ARG. The small molecules showed great potential in targeting L-ARG, which could be used to guide the development of new drugs against leishmaniasis.

Scaling up the production of rosmarinic acid and other compounds using *E. coli* could improve the feasibility of using these compounds in animal studies and clinical trials. Additionally, engineered *E. coli* provides a large avenue to explore molecular modifications that alter donor and acceptor substrates to synthesize new potential antileishmanial agents targeting the arginase enzyme.

## 2. Materials and Methods

### 2.1. Materials

CHES (2-(cyclohexylamino) ethane-sulfonic acid) and L-arginine were purchased from Sigma-Aldrich, and the reagents for urea analysis were purchased from Quibasa (Belo Horizonte, MG, Brazil). Compounds (**1-7**) were obtained via engineered synthesis in *E. coli*^[19,20]^ at Tianjin Institute of Industrial Biotechnology. Compounds 2-amino-4,5-difluorobenzoic acid, 2-amino-4-chlorobenzoic acid, trans-4-hydroxycinnamic acid, 3-(4-hydroxyphenyl)propionic acid, and trans-3,4-dihydroxycinnamic acid were purchased from Aladdin Chemistry Co., Ltd. (Shanghai, China).

### 2.2. Bacterial strain, cultivation, and chemicals

The codon-optimized hydroxycinnamoyl-benzoyl-CoA:anthranilate-N-hydroxycinnamoyl/benzoyl transferase (HCBT) was synthesized for its optimal expression in *E. coli*. 4-Coumarate CoA ligase gene (*At4CL*) was amplified via PCR from the cDNA of *Arabidopsis thaliana*. PET28a-HCBT was constructed by inserting HCBT into PET28a using the restriction sites *Nde* I and *BamH* I; PCDFDuet-At4CL was constructed by inserting At4CL into PCDFDuet using the restriction sites *Nco* I and *BamH* I. Plasmids PET28a-HCBT & PCDFDuet-At4CL / pET28a & PCDFDuet were cotransformed into *E*. *coli* BL21 (DE3), generating the recombinant strain S1 and negative control strain S2, respectively.

The strains S1 and S2 were cultivated in liquid Luria-Bertani (LB) broth or on LB agar plates at 37 °C with 50 μg/mL kanamycin and 200 μg/mL streptomycin to maintain the plasmids. For the production of biotransformation products, 1 mL of the overnight-cultured single colonies of the engineered *E. coli* strain BLRA1 was diluted with 50 mL fresh LB medium and was incubated at 37 °C, 200 rpm. When OD600 of the culture reached 0.6-0.8, 0.1 mM isopropyl-β-D-thiogalactoside (IPTG) was added to induce recombinant protein expression at 16 °C for 12 h. Subsequently, the cells were centrifuged, washed and resuspended in 50 mL of slightly modified M9 medium (1 × M9 minimal salts, 5 mM MgSO_4_, 0.1 mM CaCl_2_, and 2% (w/v) glucose, supplemented with 1% (w/v) yeast extract). The cells were treated with different feeding substrates at a concentration of 2 mM (2-amino-5-methylbenzoic acid, 2-amino-4,5-difluorobenzoic acid, and 2-amino-4-chlorobenzoic acid as acceptors; trans-4-hydroxycinnamic acid, 3-(4-hydroxyphenyl)propionic acid, and trans-3,4-dihydroxycinnamic acid as donors). The cells were cultured at 30 °C for 48 h. All substrates were purchased from Aladdin Chemistry Co., Ltd. (Shanghai, China).

### 2.3. Extraction, isolation, and identification of biotransformation products

To extract biotransformation products from the broth, fermentation was scaled up to 500 mL. After the fermentation broths were centrifuged, the supernatants containing biotransformation products were extracted using glass columns wet-packed with the macroporous resin SP825L (100 mL; Sepabeads, Kyoto, Japan). An aliquot (200 mL) of distilled water and 80% (v/v) ethanol were sequentially loaded into the column and were eluted at a constant flow rate of 1 mL/min. The eluate of 80% (v/v) ethanol was separately condensed under reduced pressure. The residue was then dissolved in 2 mL of methanol and purified via semipreparative HPLC using a Shimadzu LC-6 AD with SPD-20A detector, equipped with a YMC-pack ODS-A column (10 × 250 mm; i.d., 5 μm; YMC, Kyoto, Japan). The flow rate was 4 mL/min, and other HPLC conditions were the same as described above. The compounds were tested using NMR and MS. ^1^H-NMR spectra were analyzed on a Bruker Avance III spectrometer at 400 MHz in CD3OD. Chemical shifts were expressed as δ (ppm), and coupling constants (*J*) were expressed as hertz (Hz). MS spectra were performed on a Bruker microQ-TOF II mass spectrometer (Bruker BioSpin, Switzerland) equipped with an electrospray ionization (ESI) interface. The purity of the compounds used to test the activity was more than 90%, and compounds were characterized by NMR spectra (**Figures S1-S6**).

#### Compound 8

^1^H NMR (CD_3_OD, 400 MHz), δ H 8.83 (d, *J* = 2.1 Hz, 1H), 8.07 (d, *J* = 8.6 Hz, 1H), 7.63 (d, *J* = 15.6 Hz, 1H), 7.50 (d, *J* = 8.6 Hz, 2H), 7.14 (dd, *J* = 8.6, 2.1 Hz, 1H), 6.83 (d, *J* = 8.6 Hz, 2H), 6.52 (d, *J* = 15.6 Hz, 1H); HRESI-MS *m/z* 316.0382 [M - H]^-^ (calcd. for C16H⊓ClNO4 316.0377).

#### Compound 9

^1^H NMR (CD_3_OD, 400 MHz), δ H 8.73 (m, 1H), 7.97 (m, 1H), 7.57 (d, *J* = 15.6 Hz, 1H), 7.09 (d, *J* = 2.0 Hz, 1H), 7.00 (dd, *J* = 8.2, 2.0 Hz, 1H), 6.81 (d, *J* = 8.2 Hz, 1H), 6.46 (d, *J* = 15.6 Hz, 1H); HRESI-MS *m/z* 334.0554 [M - H]^-^ (calcd. for C16H10F2NO5 334.0527).

#### Compound 10

^1^H NMR (CD_3_OD, 400 MHz), δ H 8.69 (d, *J* = 2.2 Hz, 1H), 8.03 (d, *J* = 8.6 Hz, 1H), 7.12 (dd, *J* = 8.6, 2.2 Hz, 1H), 7.06 (d, *J* = 8.6 Hz, 1H), 6.70 (d, *J* = 8.5 Hz, 1H), 2.94 (t, *J* = 7.6 Hz, 1H), 2.69 (t, *J* = 7.6 Hz, 1H); HRESI-MS *m/z* 318.0569 [M - H]^-^ (calcd. for C16H13ClNO4 318.0533).

### 2.4. Arginase inhibition and kinetics

Recombinant L-ARG was expressed and purified as previously described.^[8]^ A stock solution of 70 mM was prepared in DMSO and was stored at −20 °C for each compound just before the experiment. The concentration that inhibits 50% of the catalytic activity of the enzyme (IC_50_). The inhibitor concentration used varied from 200 μM to 0.05 μM; the concentrations were obtained via serial dilution in water (1:4). The kinetics were determined using three concentrations of substrate and three concentrations of inhibitor, as previously described.^[10]^ Two independent experiments were performed in triplicate with a coefficient of nonlinear regression R^2^ ≥ 0.85. Sigmoidal model (log IC_50_) was used to determine IC_50_ using the GraphPad-Prism 7 software for Windows (San Diego, CA, USA).

### 2.5. Computational details

#### Molecule preparation

The three-dimensional structures of the L-ARG competitive inhibitor, compound **9**, was built in Maestro molecular modelling environment (Maestro release 2016) and minimized using MacroModel software (MacroModel, Schrödinger, LLC, New York, NY, 2016) as reported by us^[21, 22]^ employing the OPLS-AA 2005 as force field and GB/SA model for simulating the solvent effects. PRCG method with 1,000 maximum iterations and 0.001 gradient convergence threshold was employed. Furthermore, LigPrep (LigPrep, Schrödinger, LLC, New York, NY, 2016) application was used to refine the chemical structure.

#### Protein preparation

According to our previous works^[18,23,24]^ we used the recently experimentally solved structure of the *L. mexicana* arginase (PDB ID: 4IU1)^[25]^, downloaded from PDB and imported into Maestro molecular modeling environment. The structures were submitted to the protein preparation wizard protocol implemented in Maestro suite 2016 (Protein Preparation Wizard workflow 2016)^[22, 26]^ in order to obtain a reasonable starting structure for further computational experiments.

#### Molecular docking

Docking experiments were performed by Glide (Glide, Schrödinger, LLC, New York, NY, 2016) using the ligand and the protein prepared as above mentioned with the Glide extra precision (XP) method. The energy grid was prepared using the default value of the protein atom scaling factor (1.0 Å) within a cubic box centered on the crystallized ligand nor-NOHA. As part of grid generation procedure, metal constraints for the receptor grids were also applied ^[27, 28]^. After grid generation, the ligands were docked into the enzymes considering the metal constraints. The number of poses entered to post-docking minimization was set to 100, and the Glide XP score was evaluated. The XP method was able to correctly accommodated the crystallized inhibitor into the binding site (data not shown).

#### Estimated ligand binding energy

The Prime/MM-GBSA method implemented in Prime software (Prime, Schrödinger, LLC, New York, NY, 2016) computes the change between the free and the complex state of both the ligand and the protein after energy minimization. The technique was used on the docking complex of the compound presented in this study. The software was employed to calculate the ligand binding energy (ΔG_bind_) as previously reported ^[18,23,24]^.

## 3. Results

### 3.1. Production of compounds 8-10 in E. coli via feeding experiments

New three amides (**8-10**) were biosynthesized in *E. coli* cultured at 30 °C for 48 h and supplemented with 2 mM of different combinations of feeding substrates (2-amino-5-methylbenzoic acid, 2-amino-4,5-difluorobenzoic acid, and 2-amino-4-chlorobenzoic acid as acceptors; trans-4-hydroxycinnamic acid, 3-(4-hydroxyphenyl)propionic acid and trans-3,4-dihydroxycinnamic acid as donors **Scheme 1**. The compounds were characterized using NMR and MS.

**Scheme 1.**
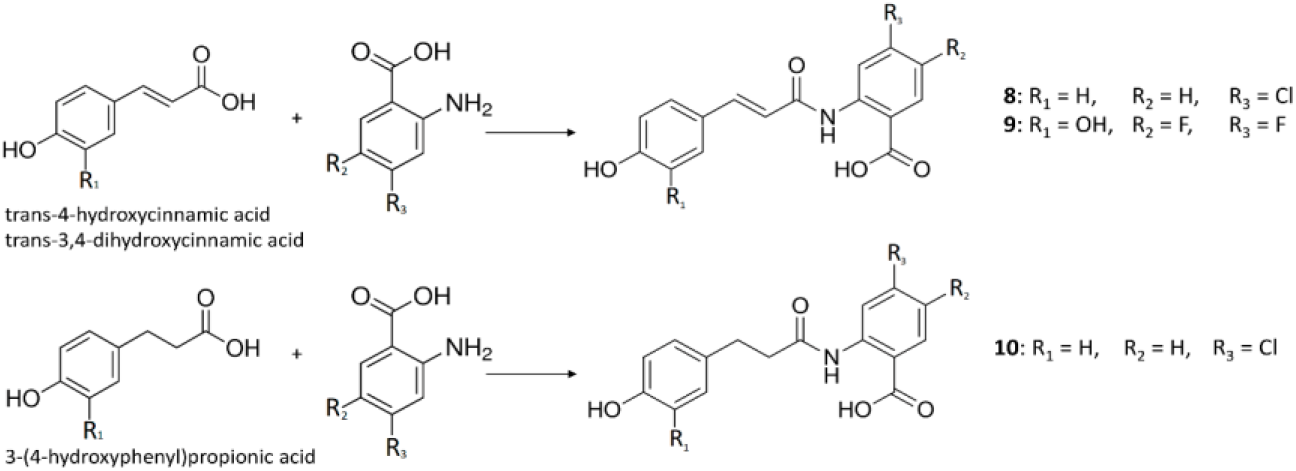
Synthesis of new cinnamids.

### 3.2. Arginase inhibition

Six biosynthetic compounds containing a catechol group, which were previously synthesized via engineered *E. coli*,^[19,20]^ showed L-ARG inhibition (**Table 1**) with IC_50_ values that ranged from 1.9 μM (**4**) to 36.2 μM (**7**). Compounds that lacked the catechol group were inactive (**1** and **6**) or showed weak inhibition (**5**) of L-ARG. The new compound **9** showed an IC_50_ value of 5.5 μM and was the unique cinnamide that was active against L-ARG (**Table 1**). Compound **9** was synthesized using 3,4-dihidroxycinnamic acid as a donor substrate and 2-amino-4,5-difluorobenzoic as an acceptor in the *E. coli* system, which generated an unnatural difluorobenzoic cinnamide with a relevant L-ARG inhibitory profile.

**Table 1.**
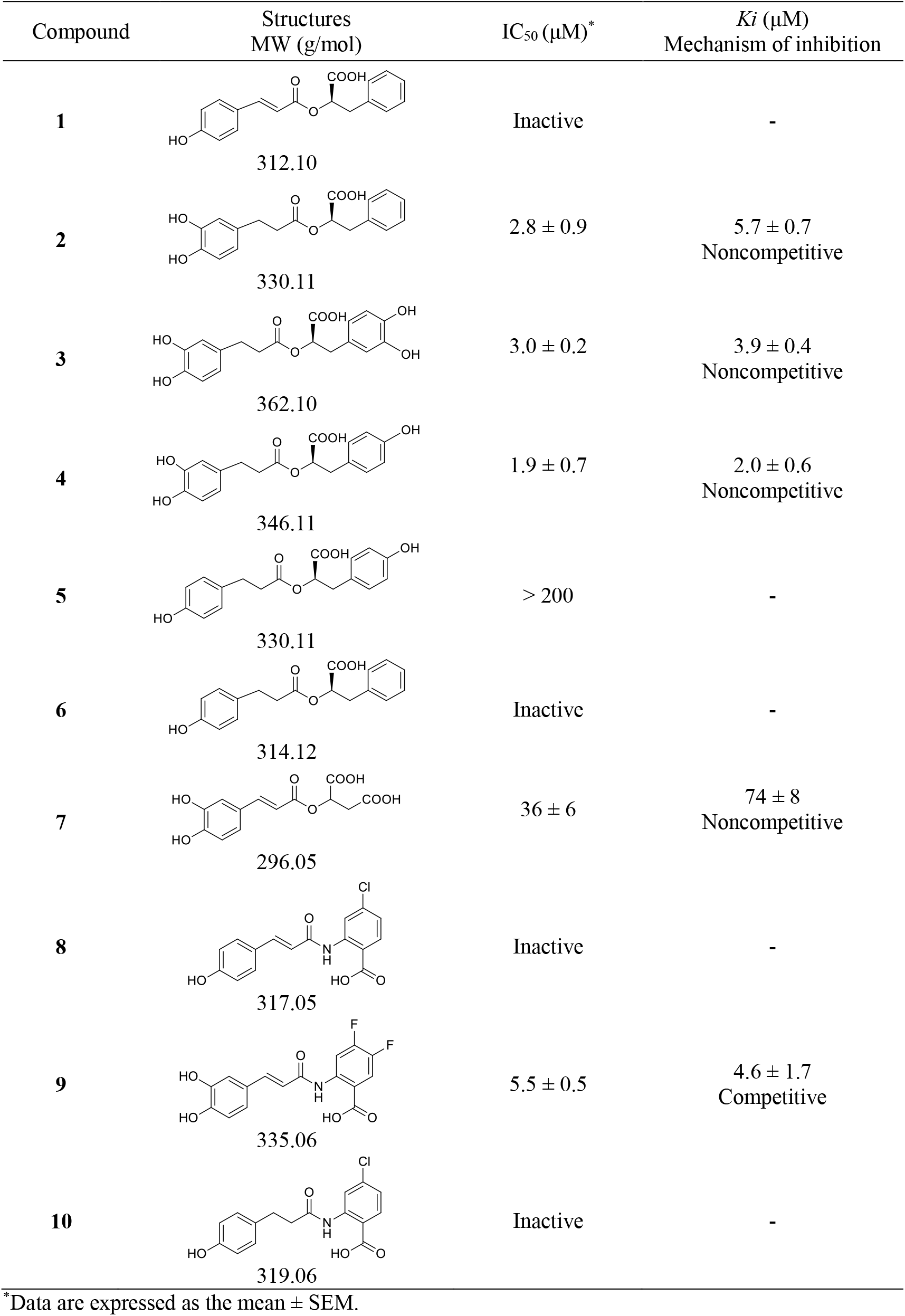
Inhibition of the *L. amazonensis* arginase by cinnamic ester compounds

| Compound | Structures<br>MW (g/mol) | IC<sub>50</sub> (μM)* | K<sub>i</sub> (μM)<br>Mechanism of inhibition |
| --- | --- | --- | --- |
| 1 | 312.10 | Inactive | - |
| 2 | 330.11 | 2.8 ± 0.9 | 5.7 ± 0.7<br>Noncompetitive |
| 3 | 362.10 | 3.0 ± 0.2 | 3.9 ± 0.4<br>Noncompetitive |
| 4 | 346.11 | 1.9 ± 0.7 | 2.0 ± 0.6<br>Noncompetitive |
| 5 | 330.11 | > 200 | - |
| 6 | 314.12 | Inactive | - |
| 7 | 296.05 | 36 ± 6 | 74 ± 8<br>Noncompetitive |
| 8 | 317.05 | Inactive | - |
| 9 | 335.06 | 5.5 ± 0.5 | 4.6 ± 1.7<br>Competitive |
| 10 | 319.06 | Inactive | - |

\*Data are expressed as the mean ± SEM.

### 3.3. Kinetics of arginase inhibition

Enzyme kinetics were performed to determine the mechanism of inhibition and determine the dissociation constant (Ki) of the six compounds via simultaneous analysis of the Dixon^[29]^ and Cornish-Bowden^[30]^ plots. The inhibition constants Ki (Complex EI) which refer to the equilibriums established between the enzyme (E) and substrate (S) in the presence of an inhibitor (I), were determined (**Table 1**). Rosmarinic acid analogs (**2-4**) showed a noncompetitive mechanism regarding L-ARG inhibition. All slopes were significantly different (p ≤ 0.05), and the interception point was used to calculate the Ki values and to obtain the mechanism of inhibition via the graph method; this result is in agreement with previous results of invariable IC_50_ obtained for three concentrations of the substrates (**Figure 1**). The new compound **9** was found to be a competitive L-ARG inhibitor (**Table 1**).

**Figure 1.**
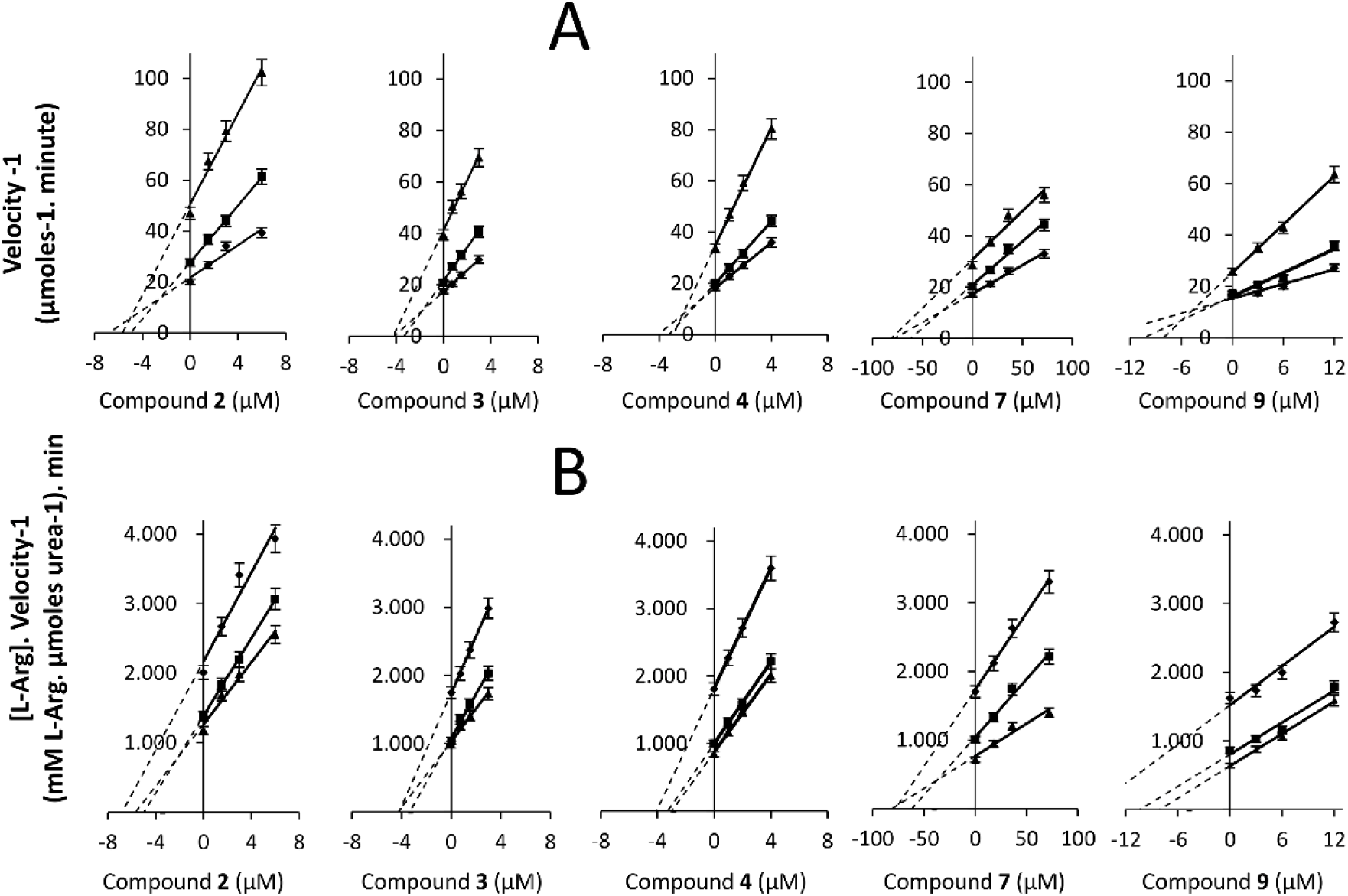
Kinetics of the arginase inhibition by compounds **2-4, 7 and 9.** The mechanism of action was determined by analysis of the Dixon (**A**) and Cornish-Bowden plots (**B**), and the *K*_i_ constant was measured using Dixon plots by calculating the X values at the intersection point of the lines. The concentrations of L-arginine were 100 mM (◆), 50 mM (■) and 25 mM (▲). Each drawn point represents the mean of three independent experiments (n = 3) performed in duplicate.

### 3.4. Computational studies

The binding mode of competitive inhibitor compound **9** was investigated applying a computational protocol based on molecular docking simulation and the estimation of ligand binding energy as previously reported by us.^[18,23,24]^ In general, we preferred to investigate only the binding of competitive inhibitors of L-ARG, since for the other inhibitors the mechanism is not completely understood and a discussion about the binding of these compounds to L-ARG could be too much speculative. Accordingly, we have performed the computational studies, employing Glide (Glide, Schrödinger, LLC, New York, NY, 2016) and Prime (Prime, Schrödinger, LLC, New York, NY, 2016) software, focused on the substrate binding site, using a crystal structure of L-ARG belonging to *Leishmania mexicana* (PDB ID 4IU1). The output of this calculation is reported in **Figure 2**.

**Figure 2.**
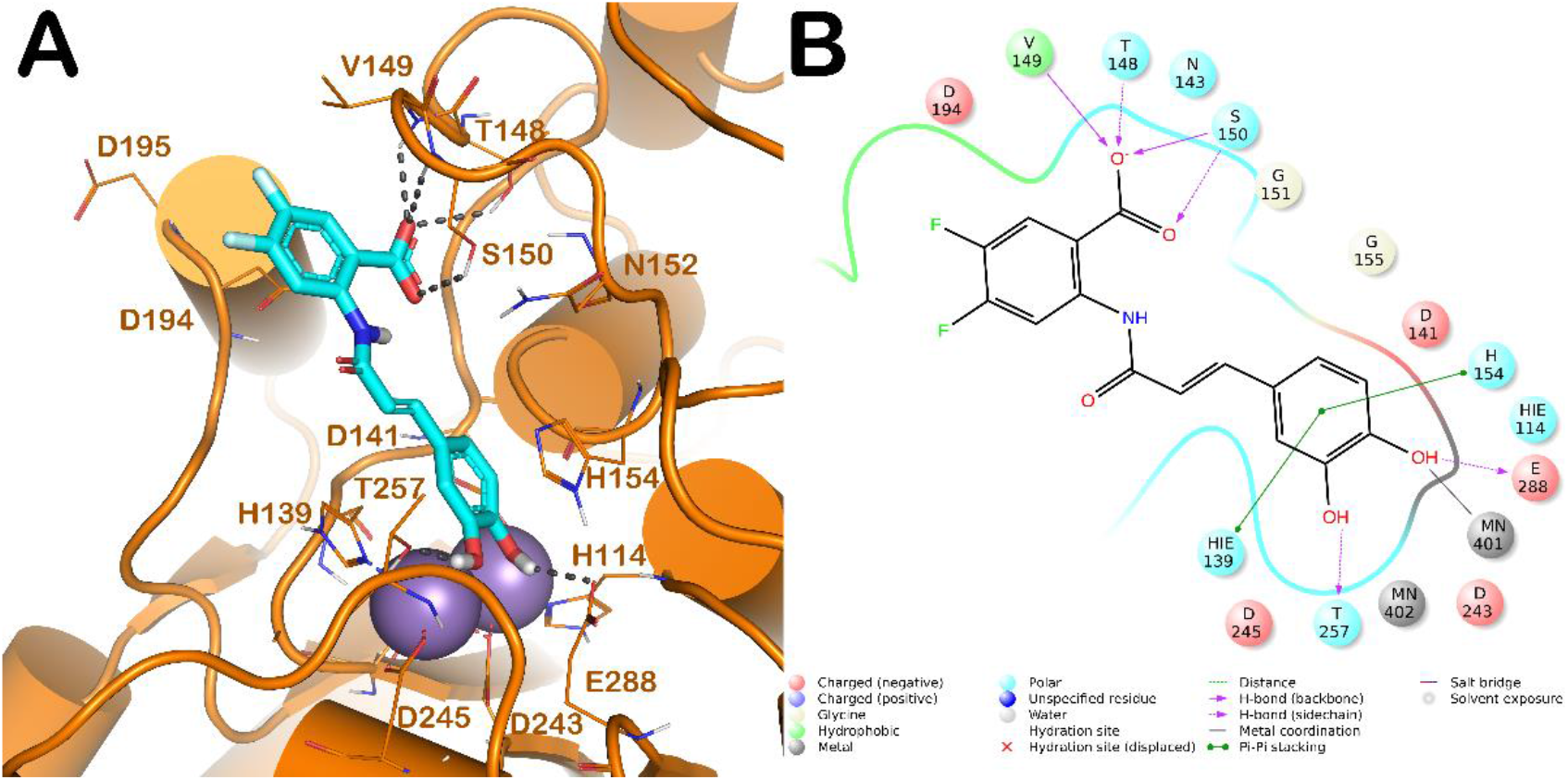
(A) Putative binding mode of compound **9** (cyan sticks) into L-ARG binding site (PDB ID 4IU1; orange cartoon) as found by Glide software (Glide, Schrödinger, LLC, New York, NY, 2016). Metals are represented as gray spheres. Key residues of the binding site are represented by lines. H-bonds are represented as black dotted lines, while the metal coordination bonds are represented by colored dotted lines. The picture was generated by means of PyMOL (The PyMOL Molecular Graphics System, v1.8; Schrödinger, LLC, New York, 2015). (B) Ligand interaction diagram (Maestro, Schrödinger, LLC, New York, NY, 2016).

Compound **9** is able to interact with L-ARG binding site by a series of polar and hydrophobic interactions. In particular, the catechol moiety is involved in a metal coordination bond by one of its hydroxyl groups. Moreover, the hydroxyl groups are able to form two H-bonds with the side chains of T257 and E288. Furthermore, the aromatic component of the catechol moiety established a double π-π stacking with H139 and H154. In addition, the carboxylic function belonging to the difluorobenzoate moiety could be able to form a strong H-bonds network by interacting with T148 (side chain), V149 (backbone) and S150 (side chain and backbone). This pattern of interaction accounted for a docking score and an estimated ligand binding energy (XP score = −8.51 kcal/mol; ΔG_bind_ = - 78.29 kcal/mol, respectively) in line with our previous computational studies performed on comparable compounds, and in line with the micromolar inhibition of the enzyme.

## 4. Discussion

Some insight could be highlighted via analysis of the six compounds that were active against L-ARG, and the results were compared with those obtained using a previous model of mammalian arginase.^[31]^ Donor substrates that contained catechol were substituted with a donor containing a phenyl moiety (trans-4-dihydroxycinnamic acid or 3-(4-hydroxyphenyl)propionic acid) in **1, 5**, and **6**; for these compounds, the inhibition of the enzyme decreased by at least 100 times (compound **4** vs. **5**) or resulted in inactive compounds (**1** and **6**). These data enhanced the importance of the 3,4-dihydroxyphenyl moiety in a donor substrate to synthesize *L. amazonensis* arginase inhibitors. Compounds **2**-**4** were synthesized using a common donor substrate (3,4-dihydroxyphenylpropanoic acid) and by varying the acceptor (4-hydroxyphenyllactic acid (HPL), phenyllatic acid (PL) and 3,4-dihidroxyphenyllactic acid (DHPL)); the analysis of L-ARG inhibition by these compounds showed that the best arginase inhibition was obtained when the acceptor used was HPL. *Leishmania* regulated the gene expression of cationic transporters and increased the uptake of L-arginine^[32]^; this could be a possible resistance mechanism if the mixed inhibitor of arginase is used as an antileishmanial agent. Therefore, as an antileishmanial drug candidate, a noncompetitive inhibitor would be a better arginase inhibitor. The replacement of hydroxyphenyl from **4** (IC_50_ 1.9 μM) to the carboxylate moiety in **7** (IC_50_ 36.2 μM) decreases the IC_50_ by approximately 20 times. Finally, compounds that were obtained using the same donor (trans-3,4-dihydroxycinnamic acid) showed that the difluorobenzoic moiety in the acceptor, which was used to obtain the cinnamide (**9**), exhibits better activity than the succinic moiety (**7**), which was used for L-ARG inhibition.

Recently, the study of natural compounds isolated from *Pluchea carolinensis*^[17]^ showed that caffeic acid (3,4-dihydroxycinnamic) and its derivative compounds chlorogenic acid, ferulic acid, and rosmarinic acid are active against promastigotes and amastigotes of *L. amazonensis*. Ferulic, rosmarinic and caffeic acids are effective in reducing the lesion size and parasite burden in the BALB/c mice model of cutaneous leishmaniasis with *L. amazonensis*.^[17]^

*In silico* investigation employing a computational protocol based on molecular docking coupled to the ligand binding energy estimation has been demonstrated by us to be a useful approach for a reliable prediction of the putative binding modes and affinity of the competitive inhibitors for L-ARG. In this case for compound **9**, acting as a competitive inhibitor of the enzyme, we observed its ability to target key residues within the binding site of the enzyme. The catechol moiety is projected toward the active site metal ions and the compound is able to establish a metal coordination bond. In addition, with the same moiety, compound **9** is able to target two additional residues in the reactive metal pocket. The other portion of the molecule by a strong H-bonds network can stabilize the retrieved binding mode interacting with a series of residues located at the entrance of the catalytic cleft. This binding mode of compound **9** supported its low micromolar inhibition of the L-ARG.

## Conclusions

This new biosynthesis approach increases the possibility of producing L-ARG inhibitors via engineered *E. coli* and opens the possibility of exploring unnatural substrates to produce new compounds with the clinical potential to treat leishmaniasis. Specifically, cinnamic esters and cinnamide-derived compounds reveal an approach to design other L-ARG inhibitors.

## ACKNOWLEDGMENTS

This research was supported by grants #17/06917-4 #19/23769-4, São Paulo Research Foundation (FAPESP), and grant #31770104, National Natural Science Foundation of China (NSFC). ERS is recipient of research productivity fellowships from the CNPq (#308218/2017-5) and JAASSC is a fellowship from Ministério da Ciência e Tecnologia Ensino Superior e Técnico-Profissional de Moçambique (MCTESTP).

## Notes

### Competing Interest Statement

The authors have declared no competing interest.

