## Supplementary material for "New cinnamid and rosmarinic acid derived compounds biosynthesized in *Escherichia coli* as *Leishmania amazonensis* arginase inhibitors": Figures S1-S6

[a] Programa de Pós-graduação em Biociência Animal, Faculdade de Zootecnia e Engenharia de alimentos, Universidade de São Paulo, 13635-Pirassununga, SP, Brazil.

[b] Departamento de Pré-Clínicas, Faculdade de Veterinária, Universidade Eduardo Mondlane, Av. de Moçambique, Km 1.5, CP 257, Maputo, Moçambique.

[c] Tianjin Institute of Industrial Biotechnology, Chinese Academy of Sciences, Tianjin 300308, China

[c] Key Laboratory of Systems Microbial Biotechnology, Chinese Academy of Sciences, Tianjin 300308, China

[d] Laboratório de Farmacologia e Bioquímica (LFBq), Departamento de Medicina Veterinária, Faculdade de Zootecnia e Engenharia de Alimentos, Universidade de São Paulo, Pirassununga, SP, 13635-900, Brazil.

**These authorscontribute equally.*

Correspondence:
**Edson R Silva,** Faculdade de Zootecnia e Engenharia de Alimentos, Universidade de São Paulo, Pirassununga, SP, 13635-900, phone: +55-19-35656828,
**Tao Liu**, Tianjin Institute of Industrial Biotechnology, Chinese Academy of Sciences, Tianjin 300308, China phone: +86-22-24828718

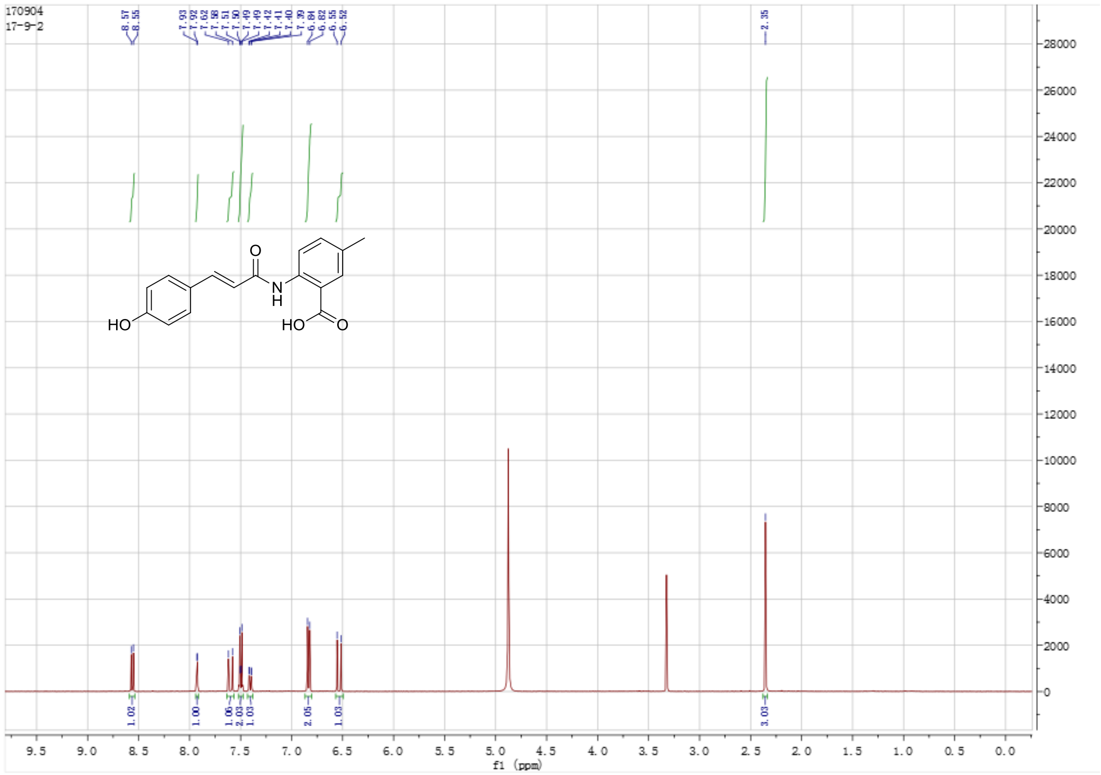


**Supplementary Figure 1**. $^{1}$H spectrum of isolated **Compound 9** in $CD_3OD$.

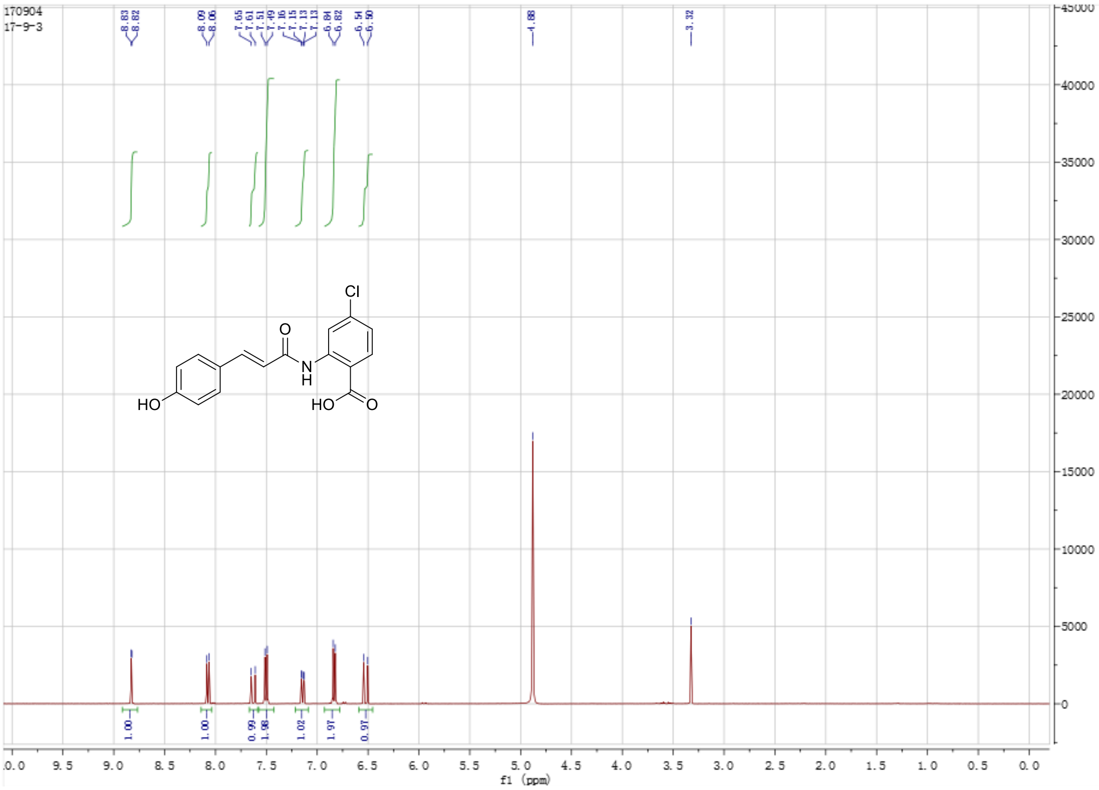


**Supplementary Figure 2**. $^{1}$H spectrum of isolated **Compound 10** in $CD_3OD$.

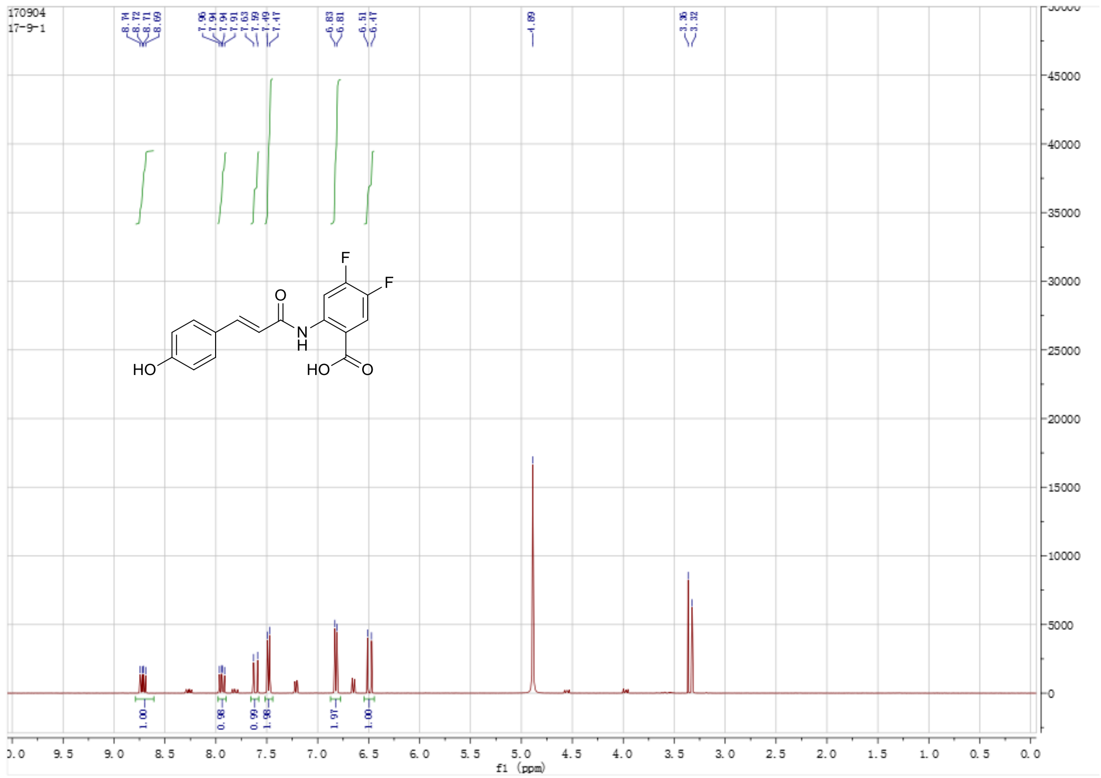


**Supplementary Figure 3**. [1]H spectrum of isolated **Compound 11** in $CD_3OD$.

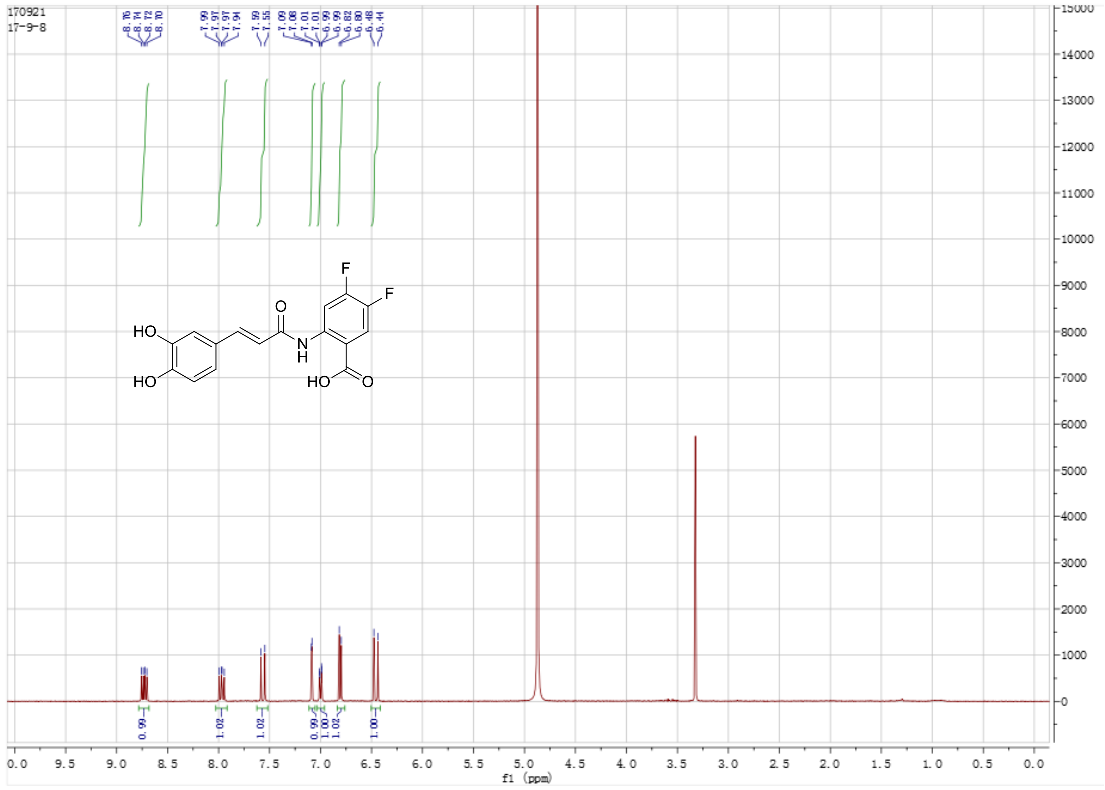

**Supplementary Figure 4**. [1]H spectrum of isolated **Compound 12** in $CD_3OD$.

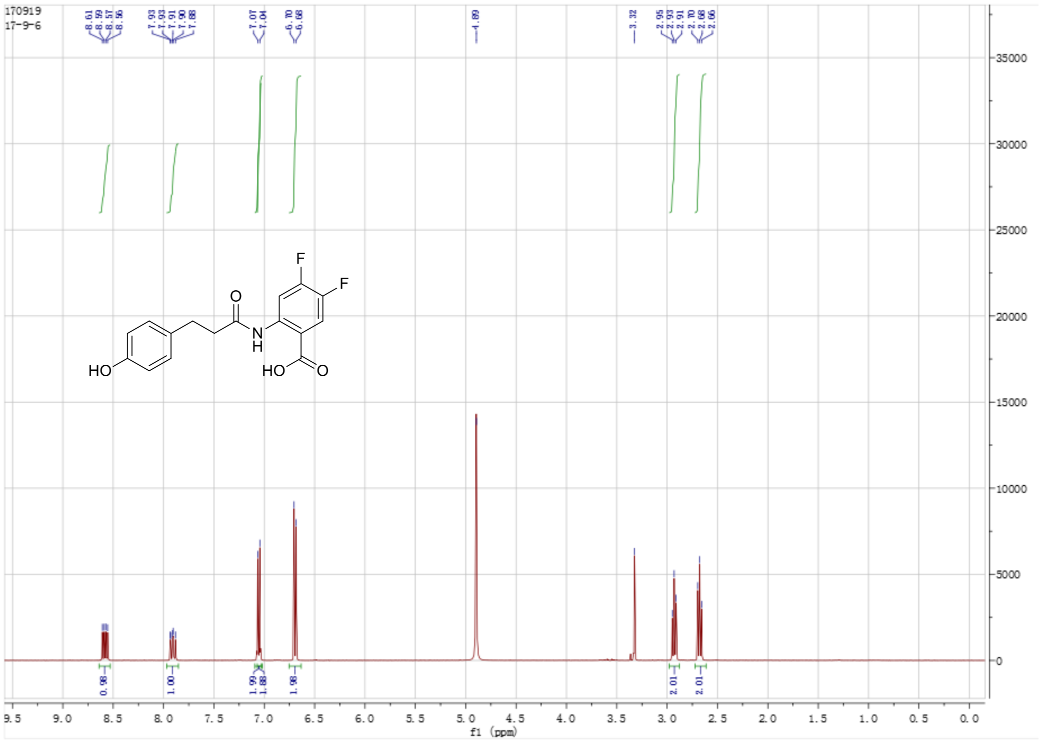


**Supplementary Figure 5**. [1]H spectrum of isolated **Compound 13** in $CD_3OD$.

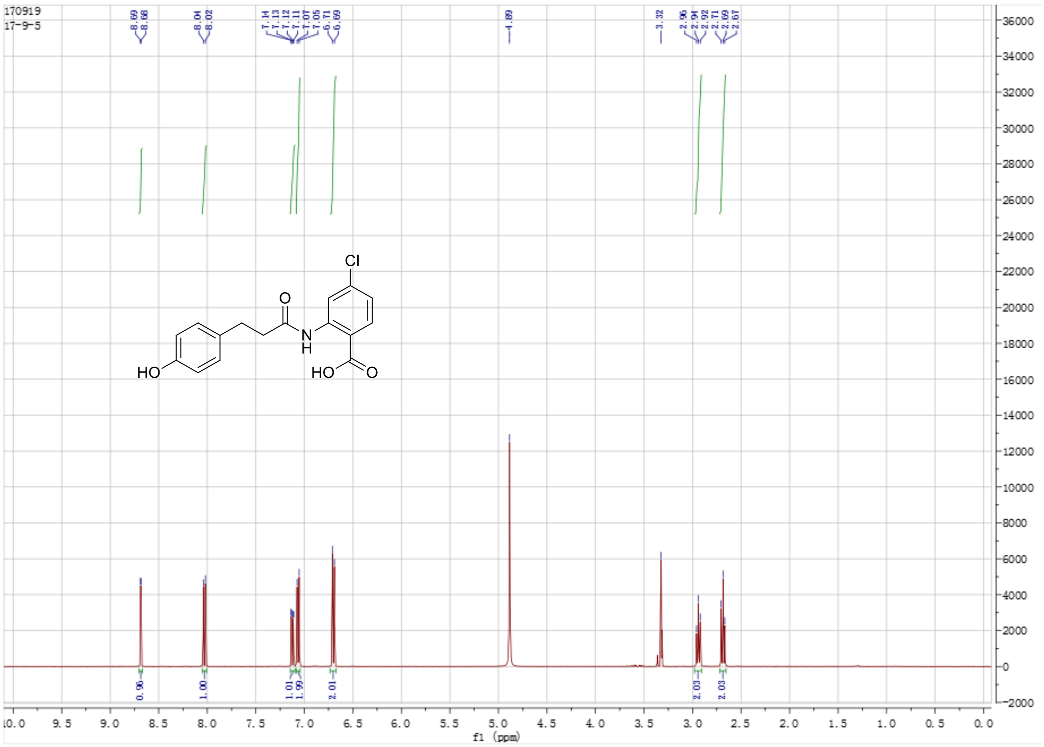


**Supplementary Figure 6**. [1]H spectrum of isolated **Compound 14** in $CD_3OD$.
